# Targeting Ubiquitin-Specific Protease 7 (USP7): A Pharmacophore-Guided Drug Repurposing and Physics-based Molecular Simulations Study

**DOI:** 10.1101/2025.04.07.647568

**Authors:** Duaa Kanan, Tarek Kanan, Berna Dogan, Ismail Erol, Serdar Durdağı

## Abstract

Ubiquitin-Specific Protease 7 (USP7) has emerged as a critical therapeutic target in cancer due to its regulatory roles of tumor suppressors, oncoproteins, and epigenetic modifiers. In this study, we present a comprehensive drug repurposing strategy to identify potential USP7 inhibitors from a curated library of 6,654 FDA-approved and investigational small molecules. Using structure-based pharmacophore models derived from USP7-inhibitor crystal structures, we conducted virtual screening to select candidates with favorable pharmacophoric alignment. The top 100 hits were subjected to short 10 ns molecular dynamics (MD) simulations and MM/GBSA binding energy calculations, narrowing down to 36 promising ligands. These were further evaluated through longer (100 ns) MD simulations, binding energy refinement, ligand clustering based on molecular fingerprints, and cancer-specific activity predictions using a binary QSAR model. Twelve compounds demonstrated favorable binding profiles, structural diversity, and high predicted anticancer activity, such as xantifibrate, etofylline clofibrate, and carafiban. This multi-tiered hybrid virtual screening pipeline highlights the potential of drug repurposing for the rapid identification of USP7 inhibitors, offering a cost-effective path toward novel anticancer therapeutics.

## Introduction

Ubiquitin specific protease 7 (USP7) or herpesvirus-associated ubiquitin-specific protease (HAUSP) is a highly promising protein target for the discovery and development of novel cancer therapeutics.^1,2^ Due to its function as a deubiquitinating enzyme, it plays a critical role in the regulation of numerous proteins that are involved in tumor suppression, immune response, cell cycle control, and DNA repair among other cellular processes.^1,2^ Through its regulation of critical proteins and cellular pathways such as the MDM2/MDMX-p53 pathway^2,3^, USP7 is known to promote the development and progression of cancer.^3–7^ USP7 also has a role in viral-associated cancers such as EBV-related nasopharyngeal carcinoma and Burkitt’s lymphoma.^8^

Accumulating evidence shows that USP7 is overexpressed in various types of cancer including multiple myeloma, chronic lymphocytic leukemia (CLL), glioma, neuroblastoma, colorectal cancer, cervical cancer, breast cancer, ovarian cancer, lung squamous cell carcinoma and large cell carcinoma, hepatocellular carcinoma, melanoma and prostate cancer.^1–4,7,9–11^ The overexpression of USP7 correlated with tumor aggressiveness and invasion in patients with prostate cancer.^11^ USP7 overexpression also correlated with worse survival of patients with multiple myeloma^12^, lung squamous cell carcinoma and large cell carcinoma^9^ as well as cervical cancer^7^. Further, USP7 has been shown to contribute to the chemoresistance of multiple cancer types such as hepatocellular carcinoma^4^ and cervical cancer.^7^

USP7 deubiquitinates and hence stabilizes a vast array of proteins. Some of its important substrates are the tumor suppressors p53 and PTEN^10^; and the MDM2/MDMX oncoproteins which are considered negative regulators of the tumor suppressor p53.^2^ Additionally, USP7 stabilizes the histone methylase EZH2, a critical protein that is involved in malignant tumor progression and invasion.^6^ Recent evidence further showed that USP7 significantly promotes the stabilization of the BCR/ABL fusion protein in chronic myelogenous leukemia (CML) and that its overexpression promotes the survival of CML cells.^5^

Due to its diverse functions, inhibition of USP7 can lead to anti-tumor effects through several mechanisms including: (i) stabilization of tumor suppressors such as p53 and PTEN^1^; (ii) decreasing the function of oncoproteins such as MDM2; (iii) regulation of epigenetic modifiers for gene expression; and/ or (iv) overcoming tumor resistance to chemotherapy.^13^ Inhibition of USP7 leads to p53 signalling activation and MDM2 degradation.^2^ In p53-mutant and p53-null CLL, USP7 inhibition effectively led to apoptosis and growth arrest via the restoration of the nuclear localization of PTEN.^10,14^ Further, the USP7-PTEN network was found to be highly aberrant in prostate cancer, in which USP7 overexpression led to the nuclear exclusion of PTEN and hence disruption of its tumor suppressor function.^11^ In another study, knockdown of USP7 was found to inhibit tumor growth and cancer invasion and led to a substantial decrease of the levels of EZH2 in prostate cancer cell lines.^6^ In CML cells, inhibition of USP7 led to significant suppression of the BCR/ABL signalling pathway and apoptosis of the CML cells.^5^ The use of USP7 inhibitor P22077 inhibited the growth of hepatocellular carcinoma in vivo and significantly overcame the chemoresistance to doxorubicin.^4^

On the other hand, use of USP7 inhibitors is suggested to increase the sensitivity of chemoradiotherapy in a variety of cancers such as multiple myeloma, acute myeloid leukemia, CLL, and solid tumors such as breast cancer.^15^ The synergistic activity of USP7 inhibitors may allow for the use of lower doses of combination therapies, thus decreasing the level of toxicities.^16^ Use of a USP7 inhibitor was found to enhance the chemotherapeutic activity of carboplatin and doxorubicin as well as the PARP inhibitor olaparib^13^, suggesting that USP7 inhibitors could work synergistically with other agents. In bortezomib-resistant multiple myeloma, use of USP7 inhibitors in combination with bortezomib led to synergistic anti-tumor activity, successfully overcoming the bortezomib resistance.^12^ USP7 inhibitors also demonstrated synergistic anti-multiple myeloma activity when used in combination with vorinostat, lenalidomide and dexamethasone.^16^

To date, several inhibitors of USP7 have been reported in the literature, but the development of highly potent and selective drugs has been slow over the years due to several factors including the limited biological and structural data on USP7^17^ and the poor knowledge of the molecular effects of its inhibition.^18^ Recently, the crystal structure of USP7 was successfully resolved in complex with small-molecule inhibitors, opening the route for more advanced structure-based drug design and development efforts.^1,19–21^ Various studies reported on developing, optimizing, and characterizing lead ligands while providing data on their activity.^21,22^ As USP7 is a highly promising drug target for cancer, it is critically important that research continues for the discovery of USP7 inhibitors that are highly specific, selective, and potent.^1,18^

By utilizing the available data and employing integrated *in silico* structure-based, ligand-based, and fragment-based approaches, we previously reported the development of pharmacophore models for USP7.^23^ A pharmacophore model represents a set of chemical features that a potential inhibitor may possess for optimal protein-ligand affinity. Pharmacophore models can be used for high throughput drug library screening like we have demonstrated. In our previous study, we reported on the discovery of seven promising USP7 inhibitors that were successful in inhibiting the activity of USP7 in *in vitro* studies.^23^ We also concluded that the structure-based complex pharmacophore model type was most successful in generating successful lead ligands.^23^

To further investigate potential inhibitors of USP7 using our structure-based pharmacophore model, we have decided to design a drug repurposing study. Drug repurposing, also known as drug repositioning or drug rescuing, is when a drug is found to have clinical use(s) other than its initial indication.^24–27^ Whereas development of a typical single drug costs from around 160 million to 2 billion dollars and takes between 11-14 years^25,27^, drug repurposing offers significant cost-saving and time-saving advantages. Computational drug repurposing allows a systematic way to analyse extensive data about chemical structures, binding, ligand-protein energy and stability, in addition to known experimental findings.^24^ Importantly, drugs that are already approved for use for other conditions may prove useful for another indication, and can go through a faster and more efficient route than traditional ways of drug development.^25^ These drugs had already passed initial safety and efficacy evaluations and are hence more likely to succeed in phase III clinical trials than drugs first entering *in vitro* studies of the drug development and testing process.^24^ For example, one of the commonly used drugs in the market, allopurinol which is currently used in the treatment of gout, was initially indicated for cancer.^27^ Another example is rituximab which was initially indicated for various cancer types but has been repurposed for rheumatoid arthritis since 2006.^24^ Other repurposed drugs that are relevant in oncology are raloxifene approved for invasive breast cancer and thalidomide used in multiple myeloma.^24,25^

In this study, we designed a drug repurposing study for the discovery of potential USP7 inhibitors as promising antineoplastic drugs. We used a structure-based pharmacophore model of the USP7 deubiquitinating enzyme to screen drugs that have already been approved by the FDA or considered investigational/experimental drugs for the discovery of promising inhibitors against the USP7 deubiquitinating enzyme. It is worthy to note that to the best of our knowledge, this is the first study of its kind that aims to investigate repurposing of drugs that are approved or considered investigational/experimental by the FDA as potential inhibitors of USP7.

Despite the growing recognition of USP7 as a central regulator in oncogenic pathways, no USP7-targeted inhibitors have yet entered clinical use, largely due to challenges in specificity, potency, and drug-likeness of newly synthesized compounds. Given these limitations, drug repurposing offers a highly promising and efficient alternative strategy to accelerate the discovery of effective USP7 inhibitors. Repurposing FDA-approved or clinically tested compounds can significantly reduce development time, cost, and safety concerns, and key barriers in anticancer drug development. Although various computational approaches have been employed to identify USP7 inhibitors, there remains a critical lack of large-scale, integrated *in silico* studies that comprehensively explore the potential of approved and investigational drugs in targeting USP7. Herein, we present the first structure-based pharmacophore-driven drug repurposing study focused exclusively on USP7, combining high-throughput virtual screening, physics-based molecular simulations, MM/GBSA binding free energy calculations, QSAR-based anticancer activity predictions, and ligand clustering. This multi-layered computational framework enables not only the identification of novel USP7-interacting compounds with favorable binding characteristics, but also highlights clinically relevant drugs that may modulate USP7-associated pathways. By bridging structure-based design with pharmacological relevance, our study provides a robust foundation for future *in vitro* investigations and paves the way for the expedited development of USP7-targeted therapies for cancer treatment.

## Methods

### Library preparation

We downloaded 7922 small molecules from the NCGC Pharmaceutical Collection (NPC) library (https://tripod.nih.gov/npc/). This library encompasses a collection of approved and investigational drugs which can be used for high-throughput screening. The approved drugs received approval from the FDA and other authorities in the world such as Health Canada. All 7922 molecules were filtered according to molecular size and number of hydrogen bond donors and acceptors, and rotatable bonds as in our previous study.^28^ We excluded molecules which weighed greater than 1000 g/mol, and which had greater than 100 rotatable bonds or greater than 10 hydrogen bond acceptor or donors. Our library included 6654 small molecules after this filtration, which was prepared using the LigPrep module in Schrodinger.

### Structure-based pharmacophore (e-Pharmacophore) model development

We used structure-based macromolecule-ligand complex pharmacophore models of USP7 ^23^ to screen our small molecule library. This pharmacophore model is generated based on the crystal structure of a protein in complex with its ligand from our previous study.^23^ To derive our pharmacophore models, we used the USP7 structure with a potent inhibitor (PDB: 6F5H). The details of this methodology were previously discussed thoroughly in our previous article.^23^ For this purpose, we generated three different models: 5-, 6- and 7-featured. The chemical features possible in the pharmacophore models are hydrogen bond acceptors (A) and donors (D); negative (N) and positive (P) ionizable features; hydrophobic (H) features; and aromatic rings (R). We previously demonstrated that this type of pharmacophore had yielded better drug screening results in comparison to the fragment-based e-pharmacophore and ligand-based pharmacophore model types.^23^ We screened our small molecule library against each of the three models and derived the top 100 molecules which matched at least 4 chemical features based on their fitness scores. Fitness is a score that gives insights into how well aligned a ligand structurally fits to the chemical features of a pharmacophore model.^29^

### Molecular Docking and Molecular Dynamics (MD) simulations

Selected top-100 compounds were docked to the binding pocket of the USP7 (6F5H) using Glide/SP. Similar docking parameters were used with our previous paper.^23^ Top-docking poses were used in MD simulations. Short (10 ns) and long (100 ns) MD simulations were conducted using Desmond. The simulations were carried out with the following parameters: OPLS3 force field; RESPA integrator; NPT ensemble at 310 K; Nose–Hoover temperature coupling; and a constant pressure of 1.01 bar maintained via Martyna–Tobias–Klein pressure coupling.^30–34^ The simulation system was prepared using the TIP3P solvent model with 0.15 M NaCl at neutral concentration. An in-house script was employed to execute the MD simulations. Initially, 10 ns MD simulations were performed for the top 100 ligands. Based on the analysis results, a subset of ligands was selected for extended simulation. For these selected ligands, as determined by MM/GBSA scoring, 100 ns MD simulations were subsequently carried out, including a positive control molecule. Additionally, replicate 100 ns MD simulations were performed for the 12 most promising compounds identified via an integrated selection strategy, along with the reference control inhibitor.

### Selection criteria

To advance from the short to long MD simulations, we considered several findings such as the MM/GBSA system free energy scores and the class of the drugs. First, we selected the top 10 molecules (out of the 100 molecules) in terms of their MM/GBSA energy scores. Thus, we identified 10 drugs as predicted to be most stable energetically based on their MM/GBSA Gibbs free energy scores. Additionally, since USP7 is implicated in cancer development, viral infections, and inflammation^8,35–37^, we decided to investigate all discovered anticancer drugs as well as anti-viral, antibiotics and anti-inflammatory drugs among top-100 compounds. Hence, we investigated the ten most energetically stable ligands and all the anti-inflammatory, anticancer, antibiotic, and antiviral drugs we discovered.

### Molecular Mechanics, General Born Surface Area (MM/GBSA) binding free energy analysis

We used the MM/GBSA free energy calculation for 10 ns MD simulations (for 100 ligands in total) and 100 ns MD simulations (for 36 ligands in total). The Prime program in Schrodinger was used for these calculations using the OPLS3 forcefield and VSGB 2.0 solvation model.^33,38^ The respective thermodynamic equations and details of this method was previously discussed in the literature.^39^ To perform these calculations, we used 100 frames throughout the simulations. A script developed in our laboratory was used to run these calculations. The value of the MM/ GBSA free binding energy for each ligand was the average of the energy scores calculated for the two chains (chains A and B of USP7).

### Clustering

The top 36 molecules based on the MM/GBSA calculation were clustered based on their similarity. The molecules were encoded using the RDKit software with Morgan fingerprints which are the implication of extended circular fingerprints.^40^ Each fingerprint is written as bit vectors of length 2048 with radius set as 3. The similarity between fingerprints, i.e. encoded molecules, were calculated using Tanimoto coefficient.^41^ The distance matrix based on Tanimoto similarity values used Butina clustering.^42^ The cut-off value for clusters were set as 0.6.

### Binary Quantitative Structure-Activity Relationship (QSAR) analysis

The MetaCore/MetaDrug platform (https://portal.genego.com) by Clarivate Analytics® allows for the analysis of ligands in terms of a multitude of structural, biochemical, and pharmacological characteristics. These characteristics include the ligands’ reactivity, blood brain barrier (BBB) penetration, predicted therapeutic activity against a variety of medical diseases and conditions as well as their predicted toxicities. For these predictions, QSAR models are used which elucidates the predicted activity of a ligand by comparing it to other ligands/ drugs used in their models. Here, we used the MetaCore/MetaDrug platform to predict the therapeutic activity of the lead 36 ligands against cancer. The binary QSAR model used to predict the potential activity against cancer has a threshold value of greater than 0.5 for predicted active ligands. This model was developed based on 886 approved drugs and drugs in clinical investigation used as the training set and 167 drugs used as the test set (*sensitivity= 0*.*89; specificity= 0*.*83; accuracy= 0*.*8*). We additionally submitted our 36 top lead ligands for toxicity prediction including for example neurotoxicity and cardiotoxicity. A thorough explanation of this platform can be found in our previous paper.^43^

### Analysis of Per-Residue root-mean-square fluctuation (RMSF) Entropy Across Ligands

To evaluate the variability in residue flexibility induced by different ligands, Shannon entropy values were calculated for each residue based on its RMSF distribution across all ligand-bound systems. First, the RMSF values obtained from MD simulations were organized into a matrix where each row represented a residue and each column corresponded to a ligand. For each residue, the RMSF values across ligands were normalized to form a probability distribution, ensuring that the sum of values for each residue equaled to 1. Subsequently, Shannon entropy *H* was calculated using the following equation:

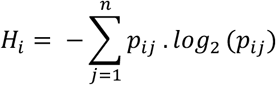

where *p*_*ij*_ is the normalized RMSF value of residue *i* for ligand *j*, and *n* is the total number of ligands. The resulting entropy values reflect the degree of variability in residue flexibility across different ligand conditions: higher entropy indicates greater variability (i.e., ligand-sensitive dynamic regions), while lower entropy denotes consistent flexibility regardless of the ligand. The entropy values were then plotted as a function of residue number to identify dynamically sensitive regions within the protein structure.

## Results and Discussion

In this study, we applied an integrated computational approach in our effort to identify potentially promising USP7 small molecule inhibitors. We screened 6654 drugs from the NPC’s NIH library, which were approved or considered experimental/investigational by the Food and Drug Administration (FDA) in the US. By using complex-based pharmacophore models of USP7 which we developed, we were able to screen thousands of molecules and assess their ability to conform to each model’s chemical features. We decided to use the structure-based/complex pharmacophore model type as we previously reported they were more successful for identifying promising small molecules than energy-optimized and ligand-based pharmacophore models.^23^ For this study, we used three different complex pharmacophore models each featuring a certain number of chemical features (5-featured, 6-featured, and 7-featured models). We screened our library against each model and identified the top 100 ligands based on their fitness scores. To further investigate our list of the potential 100 discovered ligands, these compounds were docked to the binding pocket of USP7 and top-docking poses were used in both short (10 ns) MD simulations were conducted. During the 10-ns MD simulations of 100 compounds, 100 frames per compound were recorded and average MM/GBSA calculations were performed per each compound. (Figure 1)

**Figure 1.**
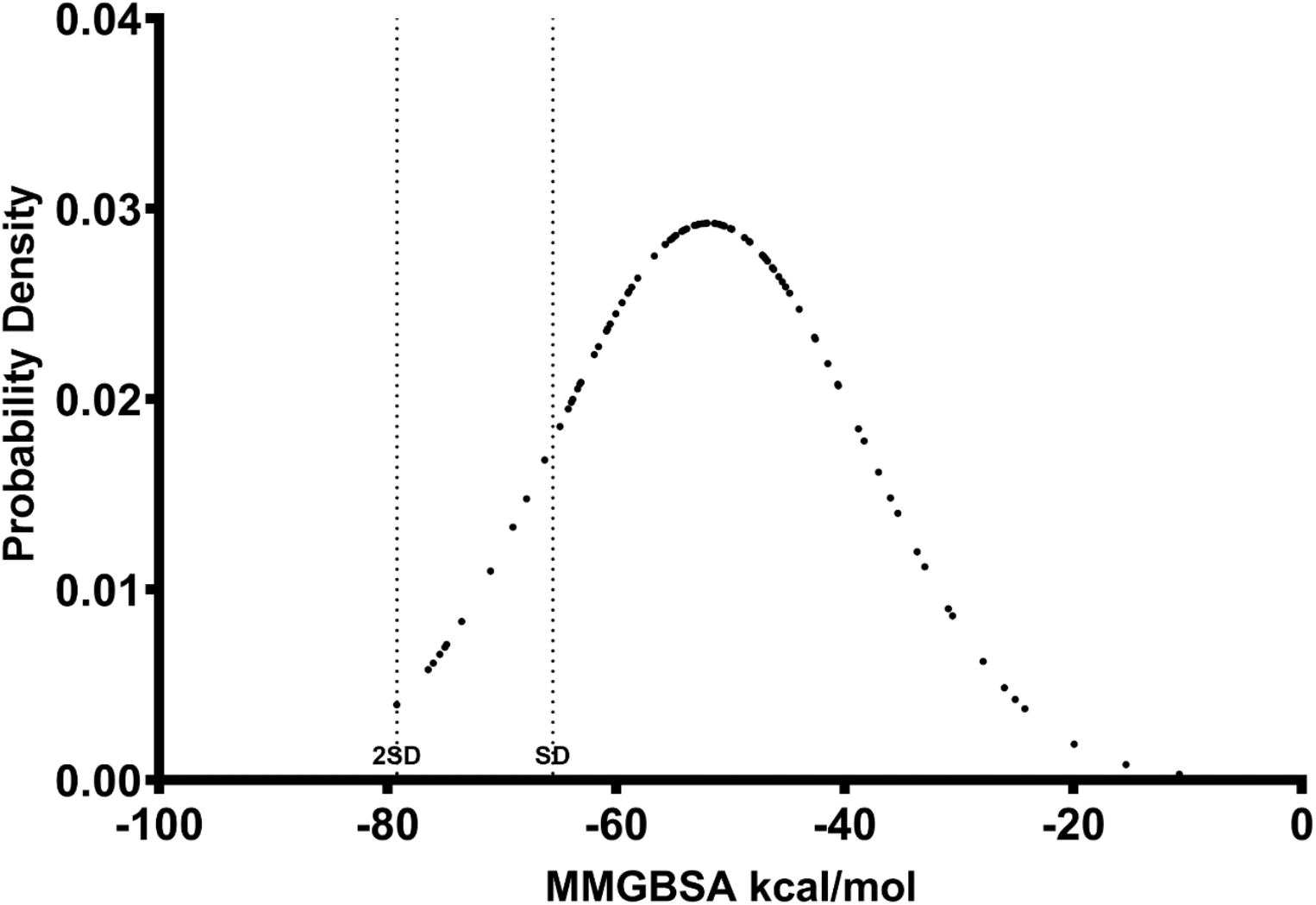
A probability density graph showing the distribution of the 100 ligands discovered via complex-based pharmacophore screening. The MM/GBSA free energy scores were calculated from the short (10 ns) MD simulations. Standard deviations (SD) away from the mean are shown.

### 100 drugs identified via e-pharmacophore screening

The 100 selected ligands were diverse in terms of function/ drug category with most drugs being identified as belonging to the class of antibiotics, antivirals and anti-inflammatory drugs followed by antilipidemics. Other drug classes included psychoanaleptics, antitussive, antihyperglycemic, thrombolytic, antihistamine, cannabinoid and anticancer drugs. It is critically important to highlight that in our search of potential anticancer drugs against USP7, we found three ligands that are already approved for use as anti-cancer drugs. These drugs were: N-(2,6-dioxo-3-piperidyl)phthalimide, also known as thalidomide; nelarabine; and fludarabine. Our findings may suggest a potential role of these drugs in interacting with and modulating its effects via USP7 or the ubiquitin-proteosome-pathway at large. To the best of our knowledge, no association between these drugs and USP7 was previously reported in the literature. Additionally, puromycin which belongs to the antibiotics drug classification has also been shown to exert cytotoxic and anti-proliferative effects in various cancer cell lines such as p53-wild type HCT116 cancer cell lines.^44^ Importantly, puromycin was shown to exert its effects via modulating the p53/MDM2 pathway and led to the reactivation/ stabilization of P53 and induction of apoptosis.^44^ Further research on analyzing the effect of puromycin on the levels of USP7, as it relates to the p53 and MDM2 pathway, will more clearly elucidate and confirm if such an association exists like proposed here in our study. Puromycin also enhanced the anti-tumor activity when used in combination with doxorubicin or the small molecule reactivating p53 and inducing tumor apoptosis (RITA), suggesting a potential role for its use synergistically in p53-wild type cancers.^44^

### MD simulations and post-MD calculations

To further investigate our list of the potential hit compounds, we extended the simulations time to 100 ns and repeated the simulations and MM/GBSA calculations. We performed 100 ns MD simulations for the top ten ligands with the lowest MM/GBSA scores ranging from –65.2 to –82.5 kcal/mol. (Table 1) The top ligands selected based on their average MM/GBSA scores (10 ligands) and which were predicted to be most potent were carafiban (a fibrinogen antagonist/ thrombolytic), etofylline clofibrate (an antilipidemic), alnespirone (an anxiolytic and antidepressant), morclofone (an antitussive), xantifibrate (an antilipidemic), droxicam (an anti-inflammatory), troglitazone (an antihyperglycemic), canbisol (a cannabinoid), barmastine (an antihistamine), and cefmatilenum (an antibiotic). Given the established role of USP7 in inflammation, pathogen-associated immune responses, viral infections, and cancer progression, we further analyzed all identified antibiotics (13 ligands), antivirals (7 ligands), anti-inflammatory agents (6 ligands), and anticancer drugs (3 ligands) among from top 100 hits.^8,35–37^

**Table 1.** The docking scores and free energy scores for the top 36 discovered hit ligands against the enzyme USP7 along with a control inhibitor. The MM/GBSA scores are provided for the 10 ns and 100 ns MD simulations separately. Drug names are listed alphabetically.

| Drug name | 100 ns -<br>MM/GBSA<br>kcal/mol | 10 ns -<br>MM/GBSA<br>kcal/mol | Docking<br>Score<br>(kcal/mol) |
| --- | --- | --- | --- |
| 1,2,3,5-Tetraacetyl-d-ribofuranose | -48.454 | -45.756 | -6.972 |
| 5-Inosinic_acid | -33.397 | -10.734 | -7.319 |
| Acetiamine | -60.035 | -42.572 | -6.385 |
| Alnespirone | -76.232 | -74.980 | -7.406 |
| Barmastine | -65.301 | -74.815 | -8.185 |
| Canbisol | -65.736 | -70.989 | -7.106 |
| Carafiban | -82.101 | -75.424 | -10.679 |
| Cefixime | -33.969 | -30.935 | -3.152 |
| Cefmatilenum | -59.317 | -53.035 | -7.826 |
| Cefmenoxime | -46.388 | -48.374 | -3.457 |
| Cefovecin | -57.197 | -63.155 | -6.914 |
| Cefpodoxime | -53.400 | -55.692 | -5.918 |
| Droxicam | -73.578 | -75.991 | -8.615 |
| Epicillin | -48.466 | -49.994 | -5.993 |
| Epiroprim | -36.990 | -49.994 | -8.445 |
| Etofylline clofibrate | -73.808 | -73.513 | -8.089 |
| Fipexide | -50.551 | -69.022 | -6.779 |
| Fludarabine | -22.982 | -32.986 | -5.284 |
| Furbucillin | -49.134 | -49.877 | -8.088 |
| Homosalate | -47.976 | -46.211 | -6.883 |
| Inosine | -54.764 | -49.877 | -7.956 |
| Isoxicam | -22.536 | -38.297 | -5.82 |
| Lodenosine | -50.888 | -26.011 | -7.967 |
| Maribavir | -51.293 | -60.741 | -5.366 |
| Metampicillin | -45.369 | -41.481 | -7.665 |
| Morclofone | -73.820 | -79.174 | -7.019 |
| Morinamidum | -47.515 | -42.647 | -6.534 |
| N-(2,6-dioxo-3-piperidyl)phthalimide | -61.542 | -63.356 | -8.051 |
| Nelarabine | -50.559 | -46.762 | -8.559 |
| Oxetacillin | -45.770 | -30.563 | -2.669 |
| Puromycin | -53.107 | -54.028 | -6.901 |
| Salnacedin | -38.534 | -50.665 | -6.497 |
| Troglitazone | -68.017 | -76.424 | -6.629 |
| valganciclovir | -45.846 | -58.102 | -8.158 |
| Vidarabine | -52.582 | -53.142 | -8.027 |
| Xantifibrate | -73.538 | -75.984 | -9.805 |
| Compound 46 | -85.881 | - | - |

Due to some overlap between the drug category of some of our top ligands based on MM/GBSA scoring with other classes, 36 lead molecules were identified in total as promising USP7 small molecule inhibitors. Thus, we performed 100 ns MD simulations for all our 36 lead molecules in addition to a known pyrimidinone-based inhibitor of USP7 (compound 46) which was derived via structure-guided lead optimization.^19^ The MM/GBSA free binding energy scores calculated from long (100 ns) MD simulations were mostly similar in both chains. Table 1 show docking scores of selected drugs and their free energy scores for all MD simulations. Figure 2 compares short and long MD simulations results.

**Figure 2.**
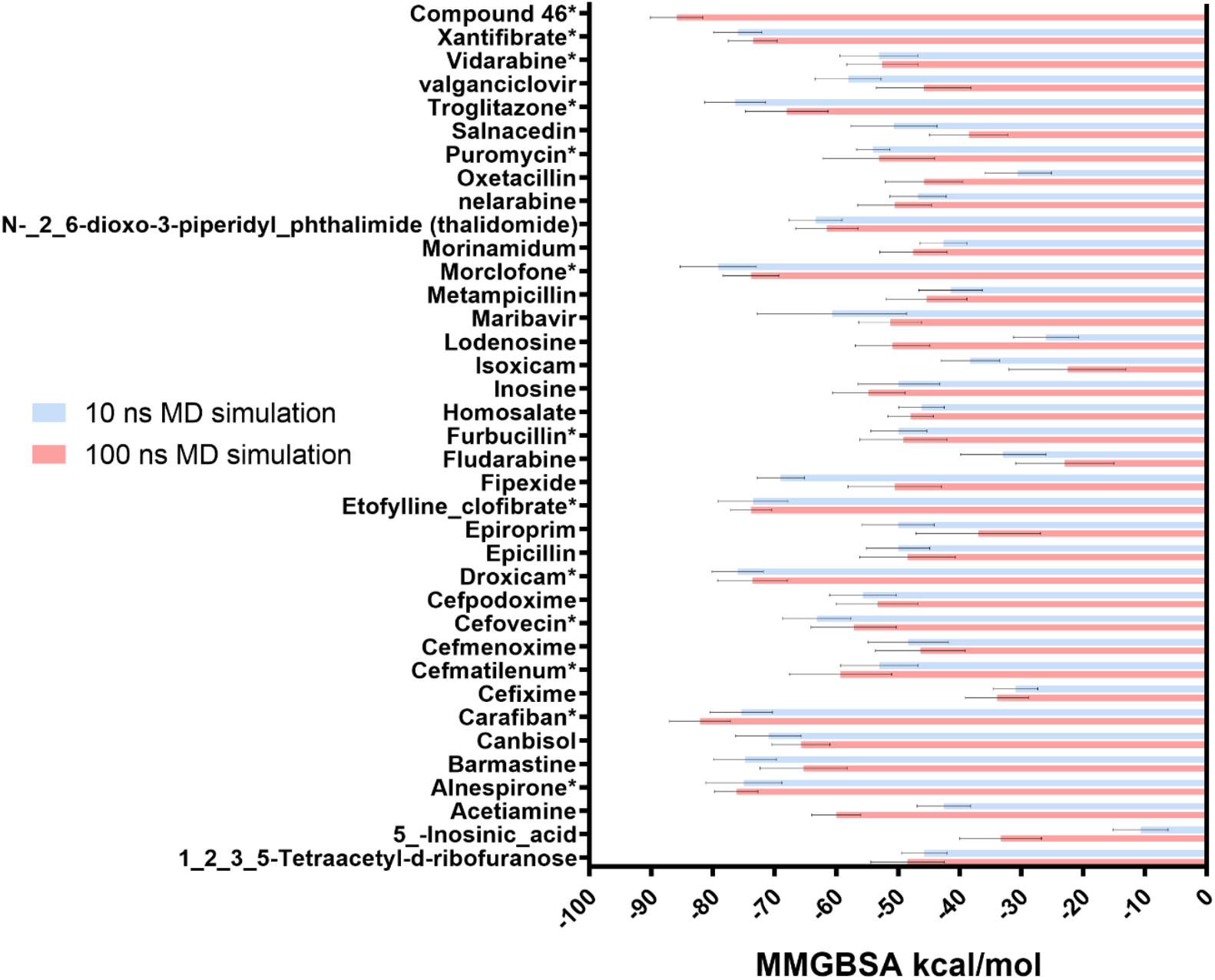
The MM/GBSA free energy scores (kcal/mol) for the 36 molecules discovered via complex-based pharmacophore screening along with a control inhibitor (Compound 46). Molecules marked with * are the identified top 12 molecules and the control inhibitor with MM/GBSA scores averaged from two replicates 100 ns MD simulations.

### Group clustering

As a result of our clustering based on Morgan 3 fingerprinting, 20 different clusters were generated. Of these clusters, only five true clusters (where a cluster is defined as a group containing more than one ligand) were identified in our study. These clusters included a variable number of ligands: cluster 1 of 8 ligands, cluster 2 of 5 ligands, cluster 3 of 4 ligands, cluster 4 of 2 ligands and cluster 5 of 2 ligands.

The Morgan 3 clustering successfully clustered ligands according to their pharmacological class: Cluster 2 ligands belong to the cephalosporin antibiotics; cluster 3 encompasses aminopenicillins such as ampicillin and amoxicillin; cluster 4 encompasses two members from the class of fenofibrates which are used clinically for hypertriglyceridemia; and cluster 5 encompasses two ligands that belong to the non-steroidal anti-inflammatory drugs (NSAIDs). This gives strong indication that ligands reported in cluster 1 (which are majorly considered nucleoside analogs) have a similar biological function considering the theme that structure dictates function.

It is important to highlight that cluster 1 includes two of the discovered anti-cancer drugs (nelarabine and fludarabine) as well as an antibiotic (puromycin) and several antiviral drugs (vidarabine, maribavir, inosine and lodenosine). The strike observation of having six ligands that are structurally similar to the anti-cancer drugs nelarabine and fludarabine, and may thus have a similar action, warrants serious investigation into assessing the function and action of these drugs in cancer and specifically in modulating USP7. The drug puromycin is being investigated for its anti-tumor effects and further research into its role shall be encouraged. Our post-MD free binding energy calculations showed that puromycin was predicted to be more potent than both nelarabine and fludarabine with an average MM/GBSA score of –53.1 kcal/mol. This cluster group also highlights the potential interaction of four different antiviral drugs in modulating USP7 and raises the question of whether the use of these drugs can be further extended to treating certain cancer types. Of the four antiviral drugs in cluster 1, inosine, nelarabine, vidarabine, maribavir and lodenosine had more favorable MM/GBSA average scores (ranging from – 50.6 to –54.8 kcal/mol) than fludarabine (–23.0 kcal/mol).

Among the identified clusters, cluster 4 had two of the top five ligands predicted to be most stable based on MM/GBSA scoring (etofylline clofibrate and xantifibrate). Our hit ligands etofylline clofibrate and xantifibrate were predicted to be highly potent USP7 small molecule inhibitors with average MM/GBSA scores of –73.8 kcal/mol and –73.5 kcal/mol respectively. A list of all ligands in each of the identified clusters is shown in Table S1.

### Analysis of the MD simulations of the top selected hit ligands

Analysis of average MM/GBSA binding free energies from extended MD simulations revealed that the top five ligands were Alnespirone, Carafiban, Etofylline clofibrate, Morclofone, and Xantifibrate. Figure 3 shows comparison analysis of these compounds with a positive control inhibitor compound 46.

**Figure 3.**
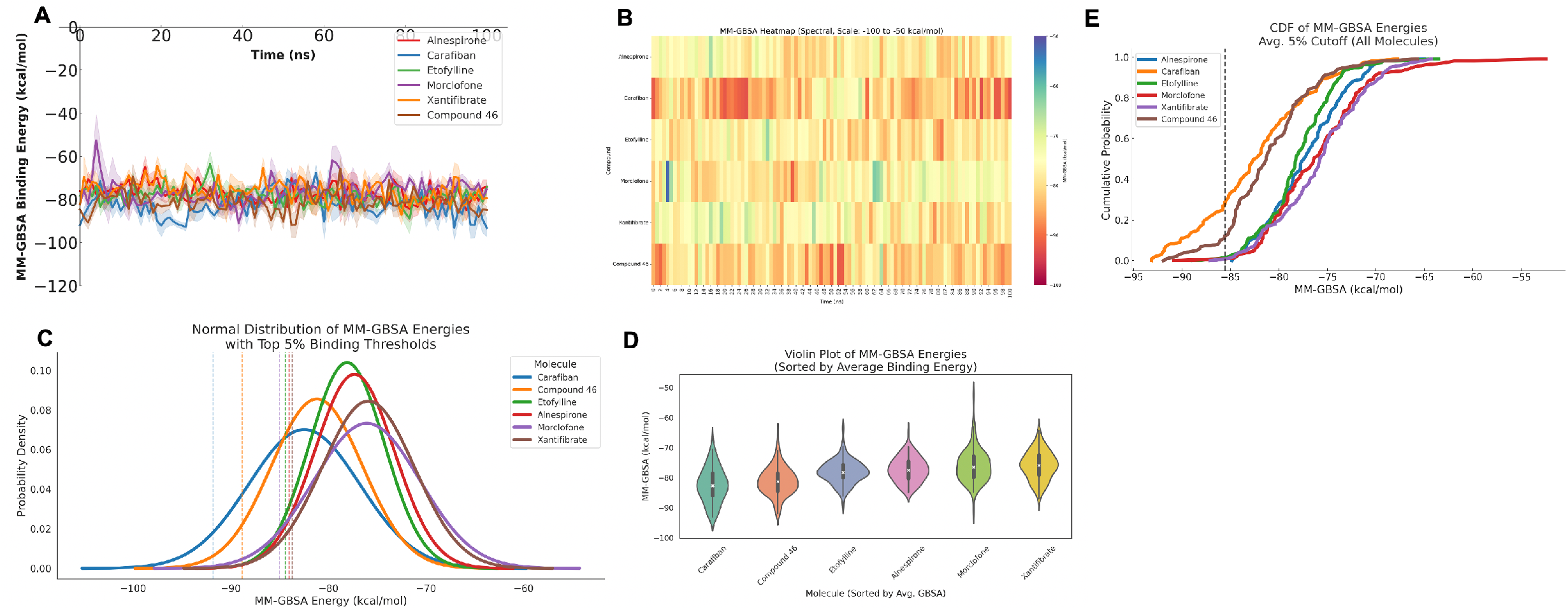
**(A)** Temporal changes in MM/GBSA binding free energies for the top five compounds and the positive control (Compound 46). **(B)** Heatmap representation of MM/GBSA binding energies. **(C)** Normal distribution plot of MM/GBSA energies with a threshold line indicating the top 5% of binding energies. **(D)** Violin plot showing the distribution and average MM/GBSA binding free energies. **(E)** Cumulative Distribution Function (CDF) of MM/GBSA values, with a dashed line representing the 5% threshold based on the average binding free energies across all compounds.

In order to determine the dynamics of the ligand-protein interactions, we have analyzed several parameters such as the root mean squared deviation (RMSD), RMSF, solvent accessible surface area (SASA) as well as the binding free energy. Considering that the RMSD of the Cα atom describes the deviation of the protein conformers when bound to ligands, our results showed that our top five ligands based on their first 100 ns MD simulation (alnespirone, carafiban, etofylline clofibrate, morclofone and xantifibrate) were relatively stable with mean Cα RMSD values of 2.1-2.3 Å compared to the control molecule with an average Cα RMSD of 1.9 Å.

Analysis of the “LigFitProtein” RMSD which helps determine fluctuations between the proteins and their ligands with respect to their initial complex alignment showed that alnespirone (RMSD 3.9 Å), etofylline clofibrate (RMSD 3.7 Å) and morclofone (RMSD 3.7 Å) had greater fluctuations than carafiban (RMSD 2.7 Å) and xantifibrate (RMSD 1.8 Å). Positive control molecule also showed some fluctuations (RMSD 2.8 Å). Hence, only one molecule – xantifibrate – is suggested to be more stable than the control molecule based on this parameter. The “LigFitLig” RMSD also provides data on the fluctuation of the ligand in comparison to its initial ligand pose in the binding pocket showed that alnespirone, etofylline clofibrate and positive control molecule had very similar fluctuations (RMSD ranging between 2.0-2.2 Å). On the other hand, carafiban, morclofone and xantifibrate demonstrated more stability when in complex with USP7 with the least fluctuation noted for xantifibrate (RMSD 1.0 Å). Hence, based on the RMSD evaluation, hit ligands carafiban and xantifibrate were particularly shown to be more stable than the positive control molecule when bound in complex with USP7. (Figure 4)

**Figure 4.**
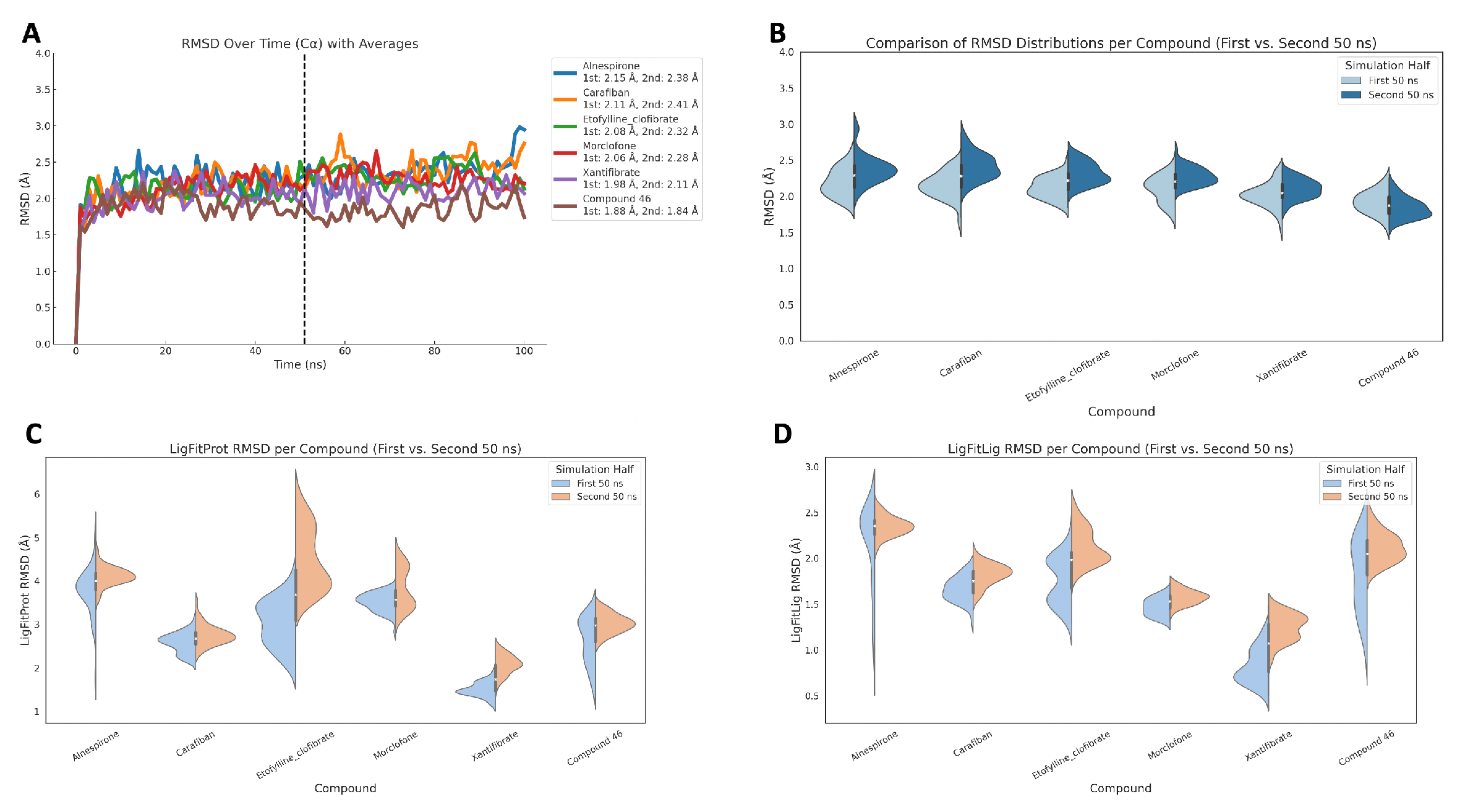
(A) RMSD of Cα atoms of the USP7 target protein plotted against simulation time. The plot compares the structural fluctuations observed during the first 50 ns and second 50 ns of MD simulations for the top five candidate compounds and a positive control ligand. (B) Distribution of protein RMSD values for each compound visualized using violin plots, summarizing the fluctuation ranges of the target protein structure across the simulation period. (C) LigFitProt RMSD distributions shown for the first and second 50 ns segments using split violin plots. (D) LigFitLig RMSD distributions across the first and second halves of the simulation time, again using split violin plots.

Another parameter which we analyzed was the RMSF Cα, which helps determine changes in the Cα position of residues in the biological system of USP7 and its ligand. All top 5 ligands (alnespirone, carafiban, etofylline clofibrate, morclofone and xantifibrate) as well as the control molecule demonstrated similar behaviour with average RMSF values ranging between 1.2-1.3 Å throughout the 100 ns MD simulations. (Figure 5)

**Figure 5.**
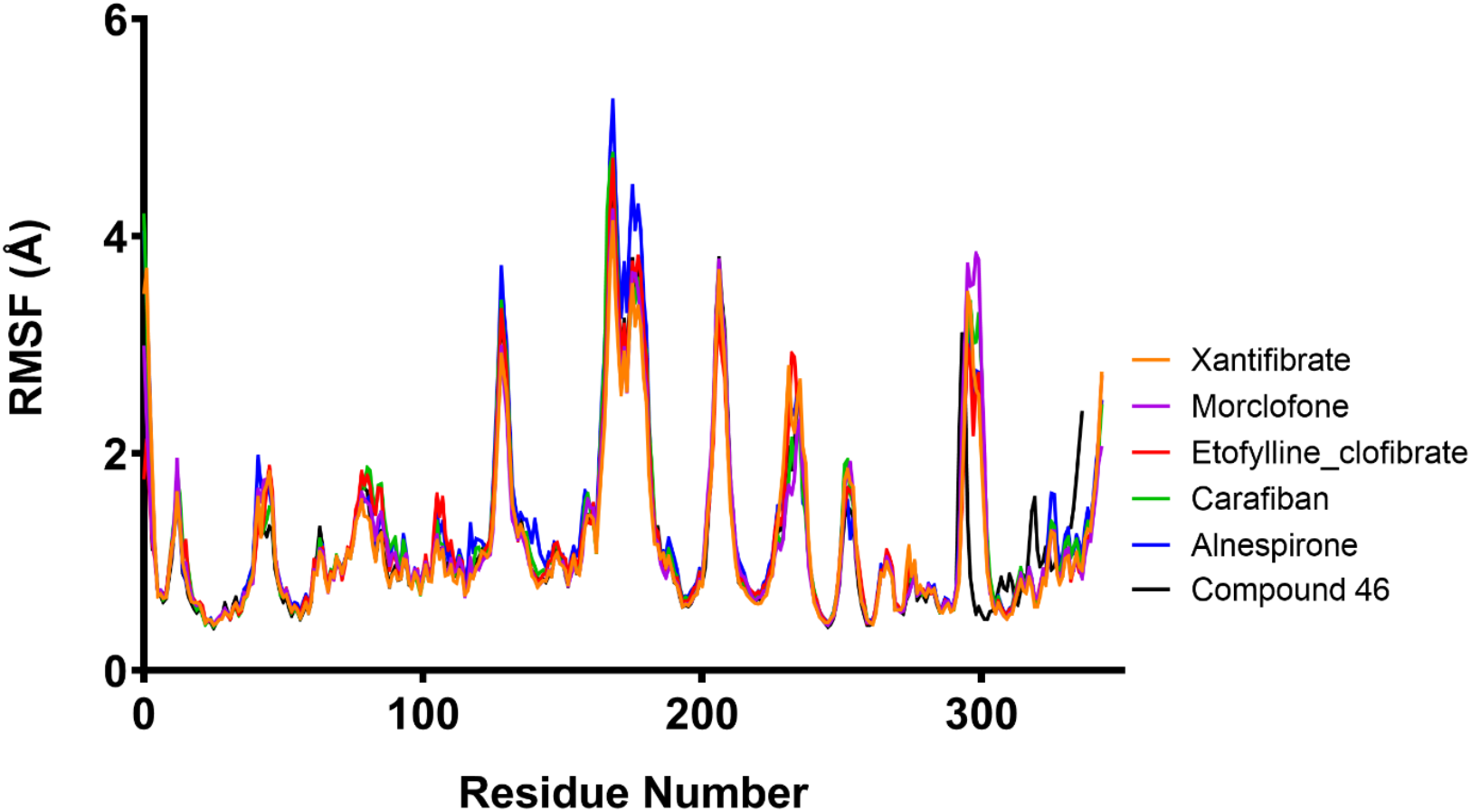
RMSF evolution of the carbon alpha atoms of the five hit ligands (based on MM/GBSA calculations) in complex with USP7 (PDB: 65FH) along with compound 46, a known USP7 inhibitor, throughout the 100 ns MD simulations.

While RMSF analysis provides insight into the average positional flexibility of residues during MD simulations, it does not account for how this flexibility varies across different ligand-bound systems. To capture this variability, Shannon entropy is applied to the per-residue RMSF profiles across all ligand complexes. In this approach, for each residue, RMSF values from all ligand-bound simulations are normalized to form a probability distribution. The Shannon entropy is then computed, quantifying the degree of variability in residue flexibility across different ligand environments. Residues with high entropy show significantly different RMSF responses depending on the ligand, indicating ligand-sensitive dynamic regions (hotspots). Conversely, low entropy residues exhibit consistent flexibility regardless of the ligand, representing structurally stable regions. This analysis allows us to distinguish between residues that are universally rigid or flexible from those whose dynamics are specifically modulated by ligand binding, providing deeper insight into allosteric behavior, ligand-specific adaptation, or potential functional sites. Figure 6 shows that specifically residues 22, 23, 26, 72 and 243 have lower entropies compared to rest of the protein. The top-entropic residues were 1, 2, 173, 174, and 214.

**Figure 6.**
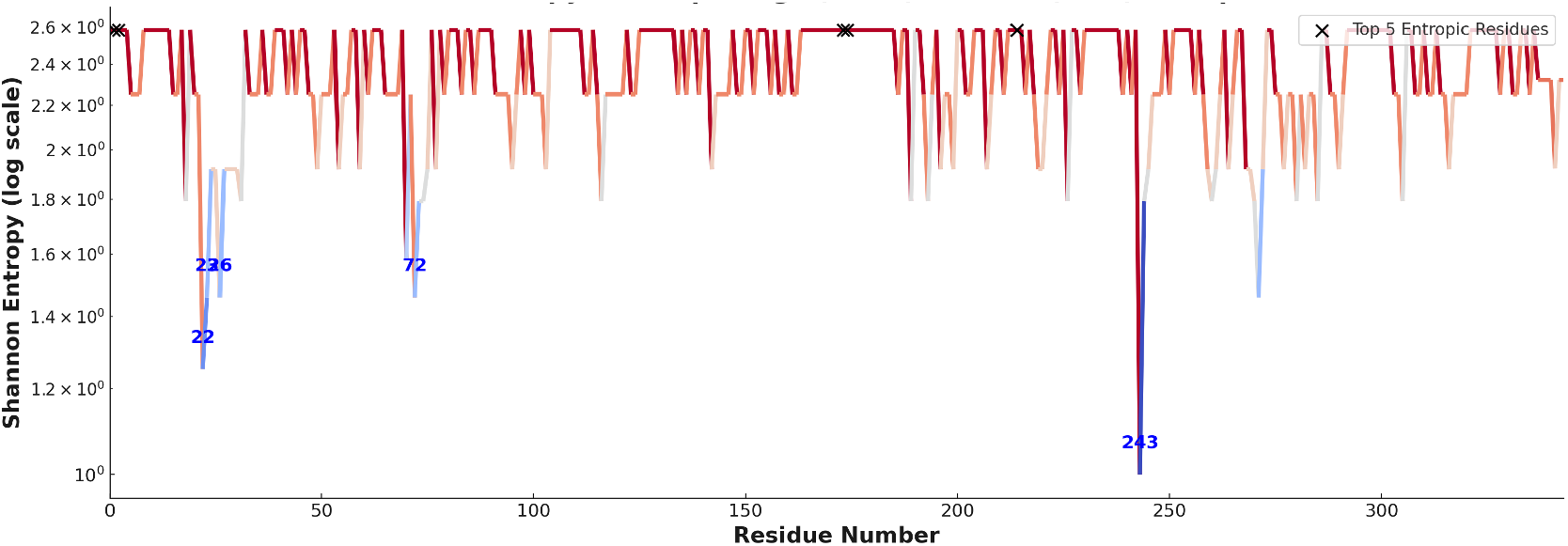
Shannon entropy values calculated per residue across top-5 hit ligands and positive control compound ligand-bound systems. The colored line plot represents the residue-wise Shannon entropy (in log scale), highlighting subtle variations in flexibility. Residues with higher entropy values exhibit greater fluctuation in flexibility across ligands, suggesting potential ligand-sensitive dynamic hotspots. The color gradient corresponds to entropy values, with red indicating higher entropy and blue indicating lower entropy.

Finally, considering solvent accessible surface area (SASA), it was shown that carafiban, etofylline clofibrate, morclofone and xantifibrate all had a lower surface area (ranging between 76.5-126.7 Å^2^) than the control inhibitor (174.3 Å^2^), likely indicating more interactions between the ligand and USP7. Only one molecule – alnespirone – had a greater accessibility surface area compared to the molecules listed above (227.2 Å^2^), Figure 7.

**Figure 7.**
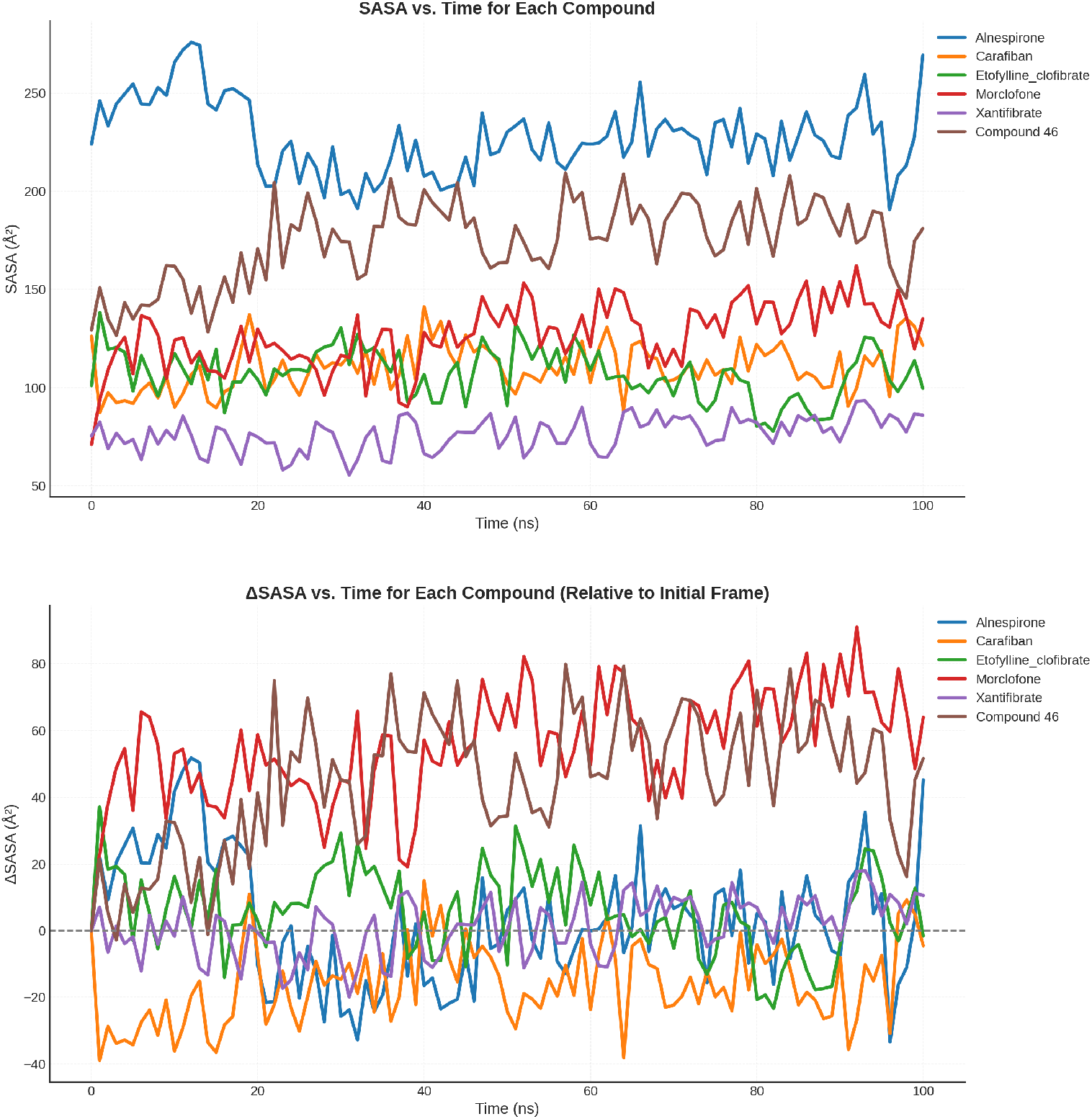
**(top)** Time-dependent changes in SASA for the top five ligands and the positive control (Compound 46) over 100 ns MD simulations. The plot illustrates how each compound modulates the exposure of the protein surface to solvent. SASA fluctuations reflect structural rearrangements and the degree of compactness or solvent exposure in the ligand-bound complexes. Notably, Alnespirone and Compound 46 exhibit higher SASA values, suggesting a more solvent-exposed or less compact complex, whereas Xantifibrate shows a relatively lower SASA profile, indicative of a more buried or compact conformation. **(bottom)** Relative changes in ΔSASA over time for the top five ligands and the positive control (Compound 46), calculated with respect to the initial simulation frame.

### MetaCore/ MetaDrug analysis of predicted activity against cancer

To further investigate selected ligands, we analyzed their predicted activity against cancer using the binary QSAR model by the Clarivate Analytics’ MetaCore/MetaDrug platform. A normalized therapeutic activity value (TAV) greater than 0.5 indicates a ligand that is predicted to be active against cancer. Of the top ten ligands we identified based on their average MM/GBSA scores, it was determined that xantifibrate (TAV = 0.82) was predicted to be most active against cancer followed by cefmatilenum (TAV = 0.74) and carafiban (TAV = 0.50) (Table S2). Xantifibrate and carafiban are also two of our top five ligands based on MM/GBSA scores as calculated from the long 100 ns MD simulations, suggesting that these two ligands are very promising small molecule inhibitors against USP7. The QSAR model used to predict the activity of molecules against cancer has successfully predicted that fludarabine and nelarabine (TAV = 0.90; TAV = 0.87, respectively) are highly active against cancer. On the other hand, N-(2,6-dioxo-3-piperidyl)phthalimide was predicted to be mildly active against cancer (TAV = 0.53). Among top 36 lead molecules, the ligand predicted to be most active against cancer was determined to be puromycin (TAV = 0.94), further adding to the power of our prediction that puromycin could have a potential mechanism of action where it modulates USP7 in order to reactivate and stabilize p53 like previously described. Further studies that analyze the effect of puromycin on the inhibition of USP7 in cancer can help elucidate whether puromycin can be considered an effective drug to be used alone or in combination with other therapeutic agents. On the other hand, another ligand which was predicted to be highly active against cancer was the anti-viral drug vidarabine (TAV = 0.90), which was identified to be in the same cluster group as fludarabine, nelarabine and puromycin.

### Integrated approach for the identification of the top 12 lead molecules

By analysis of several factors such as more favorable MM/GBSA free energy scores, calculated from the 100 ns MD simulations, predicted cancer activity as derived from the QSAR model calculations, as well as differentiation of cluster groups based on structural similarity, we identified 12 ligands as our most promising small molecule USP7 inhibitors (carafiban, alnespirone, morclofone, etofylline clofibrate, and xantifibrate, cefmatilenum, cefovecin, puromycin, troglitazone, droxicam, vidarabine and furbucillin). Among these ligands, the top five ligands with the lowest MM/GBSA scores based on the performed 100 ns MD simulations ligands were: carafiban, alnespirone, morclofone, etofylline clofibrate, and xantifibrate. Among them carafiban, etofylline clofibrate, and xantifibrate also predicted to have potential anticancer activity by cancer-QSAR model. (Figure 8) Carafiban, one of our most promising ligands, had the lowest average MM/GBSA score of –82.1 kcal/mol in comparison to the control inhibitor with an average MM/GBSA score of –85.9 kcal/mol, suggesting a highly potent small molecule USP7 inhibitor.

**Figure 8.**
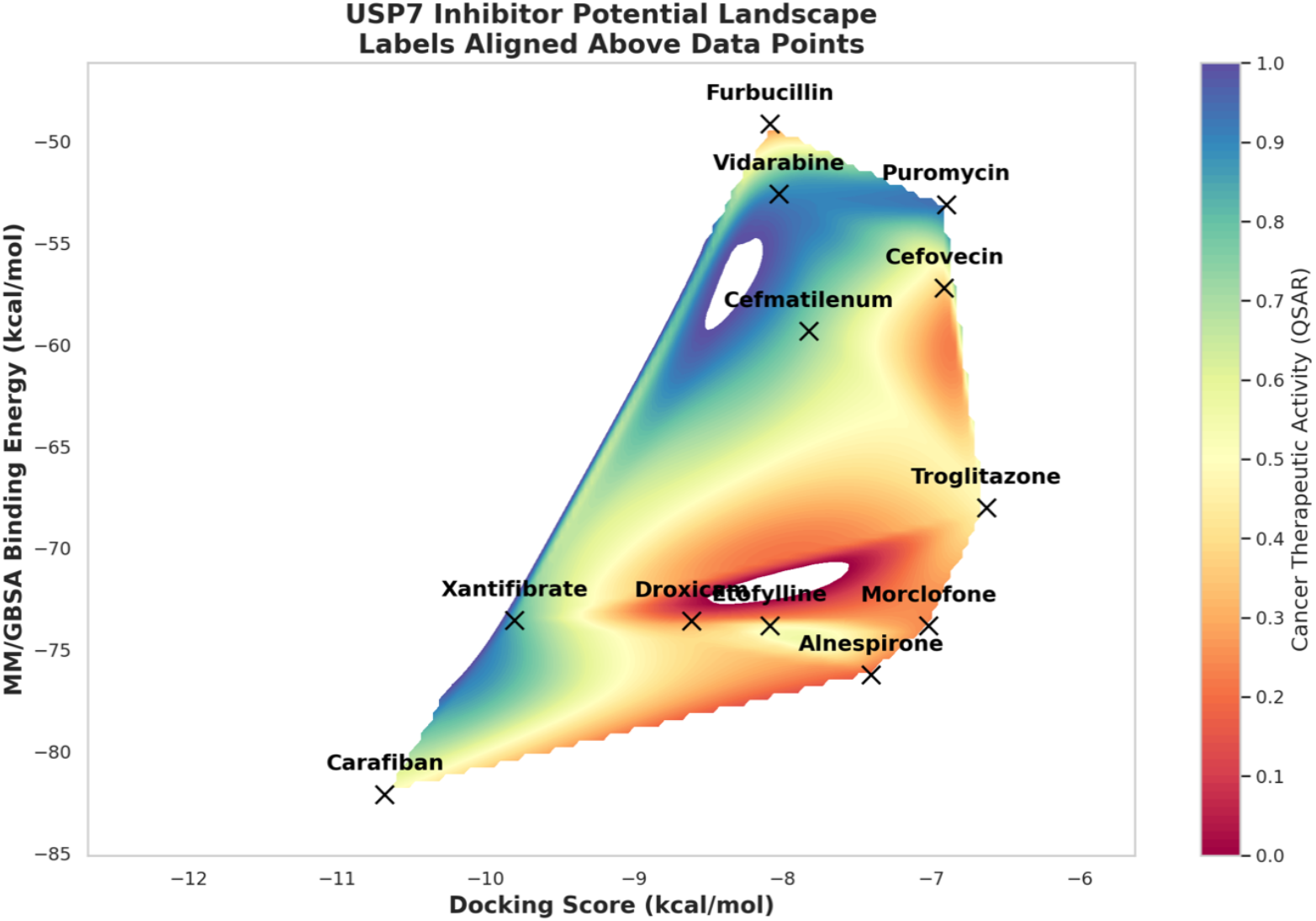
Average MM/GBSA scores, docking scores, and cancer therapeutic activity values of selected hit compounds.

## Conclusions

In this study, we employed a comprehensive pharmacophore-guided drug repurposing strategy combined with physics-based molecular simulations to identify promising small molecule inhibitors of USP7, a validated anticancer target. Through high-throughput virtual screening of a curated library of 6,654 FDA-approved and investigational compounds, 100 molecules were initially identified using a structure-based pharmacophore model derived from USP7–ligand complexes. Short and long MD simulations, along with MM/GBSA free energy calculations, enabled further refinement, leading to the selection of 36 ligands with favorable binding profiles. Among these, 12 compounds, spanning diverse pharmacological classes including antivirals, antibiotics, anti-inflammatory agents, and antilipidemics, emerged as the most promising USP7 inhibitors based on combined criteria such as MM/GBSA binding energies, QSAR-predicted anticancer activity, structural clustering, and dynamic behavior. Notably, several ligands with known anticancer properties (e.g., fludarabine, nelarabine, puromycin, and N-(2,6-dioxo-3-piperidyl)phthalimide) were identified, supporting the relevance of our computational approach and highlighting USP7 as a potential molecular target for these drugs. Puromycin, in particular, demonstrated strong binding affinity and high predicted anticancer activity, warranting further mechanistic exploration of its USP7-related actions.

Our results also revealed distinct dynamic patterns in ligand-protein interactions, as evidenced by RMSD, RMSF, SASA, and Shannon entropy analyses. These insights provided additional resolution on the ligand-induced flexibility of USP7 and helped identify ligand-sensitive dynamic regions that may serve as allosteric hotspots.

Given the complexity of USP7’s biological functions and its involvement in multiple oncogenic pathways, the rational repurposing of clinically characterized compounds presents a highly efficient strategy to accelerate USP7-targeted drug discovery. The integrated *in silico* pipeline established here, combining pharmacophore modeling, MD simulations, free energy estimations, structural clustering, and QSAR analysis offers a powerful framework for prioritizing compounds for *in vitro* validation.^46–50^

In conclusion, this study presents a robust computational foundation for the repurposing of clinically relevant compounds as potential USP7 inhibitors, paving the way for the development of novel anticancer therapeutics targeting this critical enzyme. Experimental validation of the top-ranked candidates is warranted to confirm their biological activity and therapeutic potential.

