## supporting information for "Targeting Ubiquitin-Specific Protease 7 (USP7): A Pharmacophore-Guided Drug Repurposing and Physics-based Molecular Simulations Study"

[1]Department of Internal Medicine, University of Illinois College of Medicine, Peoria, IL, 61605 USA; [2]Computational Biology and Molecular Simulations Laboratory, Department of Biophysics, School of Medicine, Bahcesehir University, 34734, Istanbul, Türkiye; [3]Department of Chemistry, Istanbul Technical University, 34467, Istanbul, Türkiye; [4]Department of Analytical Chemistry, School of Pharmacy, Bahcesehir University, 34353, Istanbul, Türkiye; [5]Molecular Therapy Lab, Department of Pharmaceutical Chemistry, School of Pharmacy, Bahcesehir University, 34353, Istanbul, Türkiye; [6]Laboratory for Innovative Drugs (Lab4IND), Computational Drug Design Center (HİTMER), Bahçeşehir University, 34734, İstanbul, Türkiye

### Supporting Tables

**Table S1.** Ligand categorization based on Morgan 3 clustering. The 36 lead ligands were analysed using clustering and Tanimoto Prioritization and members of the five clusters that were found are shown. Ligands in bold (eight) are among the top 12 ligands selected as most promising in our study based on multiple factors including ligand clustering. Ligands marked with * (two) are among the top 5 ligands based on the initial run of the 100 ns MD simulation MM/GBSA energy score analysis.

**Proposed USP7 Inhibitors As Potential Anticancer Drugs**

| Cluster 1 | Cluster 2 | Cluster 3 | Cluster 4 | Cluster 5 |
|---|---|---|---|---|
| **Vidarabine** | **Cefovecin** | **Furbucillin** | **Etofylline clofibrate*** | **Droxicam** |
| **Puromycin** | **Cefmatilenum** | Epicillin | **Xantifibrate*** | Isoxicam |
| Inosine | Cefmenoxime | Metampicillin | | |
| Lodenosine | Cefpodoxime | Oxetacillin | | |
| 5'-Inosinic acid | Cefixime | | | |
| Nelarabine | | | | |
| Fludarabine | | | | |
| Maribavir | | | | |
| ↓ | ↓ | ↓ | ↓ | ↓ |
| Nucleoside analogs | Cephalosporins | Aminopenicillins | Fenofibrates | NSAIDs |

**Table S2.** Chemical structures and characteristics of selected molecules among the top 36 discovered hit ligands against USP7. These top 12 molecules were selected based on factors such as MM/GBSA energy scores, group clustering categorization and predicted activity against cancer. Drug names are listed in alphabetical order.

| Drug name | Structure | 100 ns - MM/GBSA kcal/mol | 10 ns - MM/GBSA kcal/mol | Docking Score (kcal/mol) | Cancer therapeutic activity |
|---|---|---|---|---|---|
| Alnespirone* | 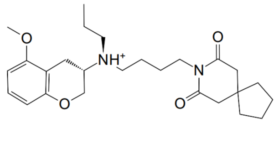 | -76.232 | -74.980 | -7.406 | 0.17 |
| Carafiban* | 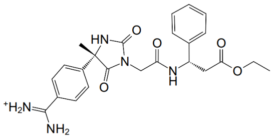 | -82.101 | -75.424 | -10.679 | 0.50 |
| Cefmatilenum | 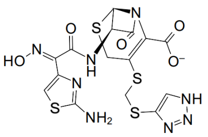 | -59.317 | -53.035 | -7.826 | 0.74 |
| Cefovecin | 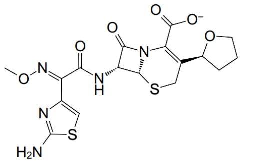 | -57.197 | -63.155 | -6.914 | 0.36 |
| Droxicam | 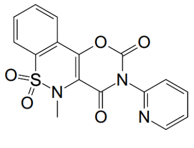 | -73.578 | -75.991 | -8.615 | 0.24 |

| | | | | | |
|---|---|---|---|---|---|
| Etofylline clofibrate* | 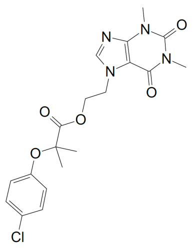 | -73.808 | -73.513 | -8.089 | 0.47 |
| Furbucillin | 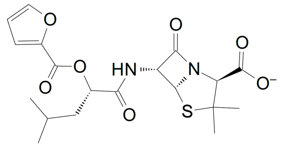 | -49.134 | -49.877 | -8.088 | 0.20 |
| Morclofone* | 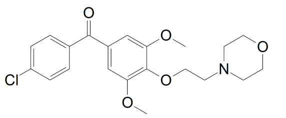 | -73.820 | -79.174 | -7.019 | 0.24 |
| Puromycin | 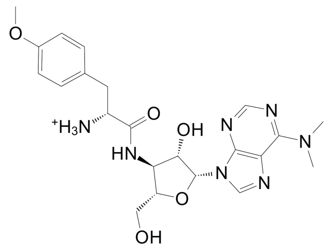 | -53.107 | -54.028 | -6.901 | 0.94 |
| Troglitazone | 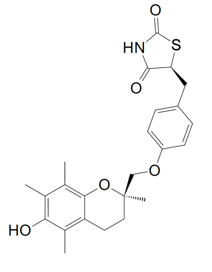 | -68.017 | -76.424 | -6.629 | 0.48 |
| | | -52.582 | -53.142 | -8.027 | 0.90 |

| | | | | | |
|---|---|---|---|---|---|
| Vidarabine | 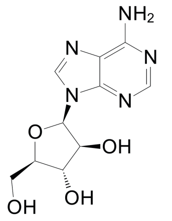 | | | | |
| Xantifibrate* | 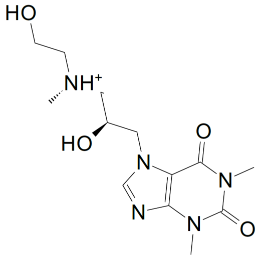 | -73.538 | -75.984 | -9.805 | 0.82 |

**Top five molecules based on their first 100 MD simulation-based MM/GBSA scores.*
